# A deep learning convolutional neural network distinguishes neuronal models of Parkinson’s disease from matched controls

**DOI:** 10.1101/2023.11.23.568499

**Authors:** Rhalena A. Thomas, Eddie Cai, Wolfgang Reintsch, Chanshaui Han, Sneha Shinde, Roxanne Larivière, Andrea Krahn, Carol X.Q. Chen, Emmanuelle Nguyen-Renou, Eric Deneault, Zhipeng You, Thomas M. Durcan, Edward A. Fon

## Abstract

Parkinson’s disease (PD) is a neurodegenerative disorder that results in the loss of dopaminergic neurons in the substantia nigra pars compacta. Despite advances in understanding PD, there is a critical need for novel therapeutics that can slow or halt its progression. Induced pluripotent stem cell (iPSC)-derived dopaminergic neurons have been used to model PD but measuring differences between PD and control cells in a robust, reproducible, and scalable manner remains a challenge. In this study, we developed a binary classifier convolutional neural network (CNN) to accurately classify microscopy images of PD models and matched control cells. We acquired images of iPSC-derived neural precursor cells (NPCs) and dopaminergic (DANs) and trained multiple CNN models comparing control cells to genetic and chemical models of PD. Our CNN accurately predicted whether control NPC cells were treated with the PD-inducing pesticide rotenone with 97.60% accuracy. We also compared control to a genetic model of PD (deletion of the Parkin gene) and found a predictive accuracy of 86.77% and 95.47% for NPC and DAN CNNs, respectively. Our cells were stained for nuclei, mitochondria, and plasma membrane, and we compared the contribution of each to the CNN’s accuracy. Using all three features together produced the best accuracy, but nuclear staining alone produced a highly predictive CNN. Our study demonstrates the power of deep learning and computer vision for analyzing complex PD-related phenotypes in DANs and suggests that these tools hold promise for identifying new targets for therapy and improving our understanding of PD.

## Introduction

Parkinson’s disease (PD) is a common neurodegenerative disorder characterized by the progressive loss of dopaminergic neurons in the substantia nigra pars compacta, affecting greater than 1% of the population aged 60 and older.(1,2) Despite advances in understanding the disease pathology, there remains a critical need for novel therapeutics that can slow or halt the progression of PD.(3) One approach to studying PD is to use human patient somatic cells reprogrammed into stem cells or induced pluripotent stem cell (iPSCs), that can be differentiated into any cell type, including dopaminergic neurons to model PD.(4) However, assessing the phenotypic differences between neurons derived from patients from those derived from healthy controls can be challenging. For instance, while it is clear that morphological differences exist between iPSC-derived PD and healthy neurons (5–9), uncovering such differences requires time-consuming and difficult-to-measure changes in microscopic images.

In addition to the challenge of accurately measuring phenotypic differences between disease and healthy control neurons, there is a critical need for scalable and high-throughput techniques to screen potential therapeutic compounds for PD. Traditional methods for drug screening, such as manual observation of cellular morphology or viability, are time-consuming and labor-intensive. Thus, there is a pressing need for efficient and reproducible methods to assess the effects of candidate drugs on cellular morphology and function.

To address these problems, we turned to the field of deep learning and convolutional neural networks (CNNs). CNNs have been widely used in image classification and segmentation tasks and have been successfully applied to biological images to identify and classify different cell types. In this study, we developed a binary classifier CNN to accurately classify microscopy images of cells modeling PD. Specifically, we used high content immunofluorescence microscopy images of organic dyes marking the nucleus, mitochondria and plasma membrane to train and test our CNN. We find that the CNN robustly and accurately classifies neural precursor cells and neurons from both a toxin and a genetic model of PD. The development of CNN-based classifiers provides a promising avenue for the rapid and reliable phenotypic screening of large compound libraries, enabling the identification of potential therapeutic candidates for PD.

## Results

### CNN models accurately predict disease status in toxin and genetic models of PD

As an initial test to assess the feasibility of using machine vision to distinguish between iPSC derived cells, we used the pesticide rotenone as an *in vitro* model of PD.(10,11) The healthy control iPSC line AIW002-02 was used for these initial studies, and was differentiated into dopaminergic neuronal precursor cells (NPCs), as previously described.(12) Half the cells were treated with rotenone then fixed and stained with organic dyes, wheat germ agglutinin (WGA) to label cell membranes and Hoechst to label the nucleus. Images of NPCs were processed and used to train a CNN model to distinguish between the rotenone treated (disease state) and untreated (healthy) NPCs.

We designed our binary CNN classifier using an adapted version of the VGG model.(13) Our model contains six convolutional layers, three max pooling layers, two dense fully connected layers and the output layer (**Figure 1A**). The processed image dataset was split randomly to remove 20% of the images as a hidden test set. The remaining 80% of the images were used to train the CNN. For training, the training image set is again split into a training set and a validation set (80/20), the validation images are not used to train the model, but rather to test the model accuracy during training. We used a batch size (number of training samples shown to the model before the model’s internal parameters are updated) of 64 and 128 steps (number of batches) per epoch. The model was trained until 15 epochs passed without a change in error rate. For each epoch of training, the error loss and accuracy of the model were calculated (**Figure 1B and C**). The goal of the CNN model is to reduce the loss function, the loss starts high and quickly drops, gradually decreasing until plateauing after about 800 epochs around 0.25 (**Figure 1B**). The model accuracy shows the inverse pattern, starting at 50% and quickly improves, leveling out at ∼90% with minor fluctuations after only about 400 epochs (**Figure 1B and C**). Next, we predicted if the hidden test images of NPCs were of cells treated with rotenone or not. We found the CNN model made minimal mistakes in predicting disease state with an accuracy of 97.60% (**Figure 1D**).

**Figure 1:**
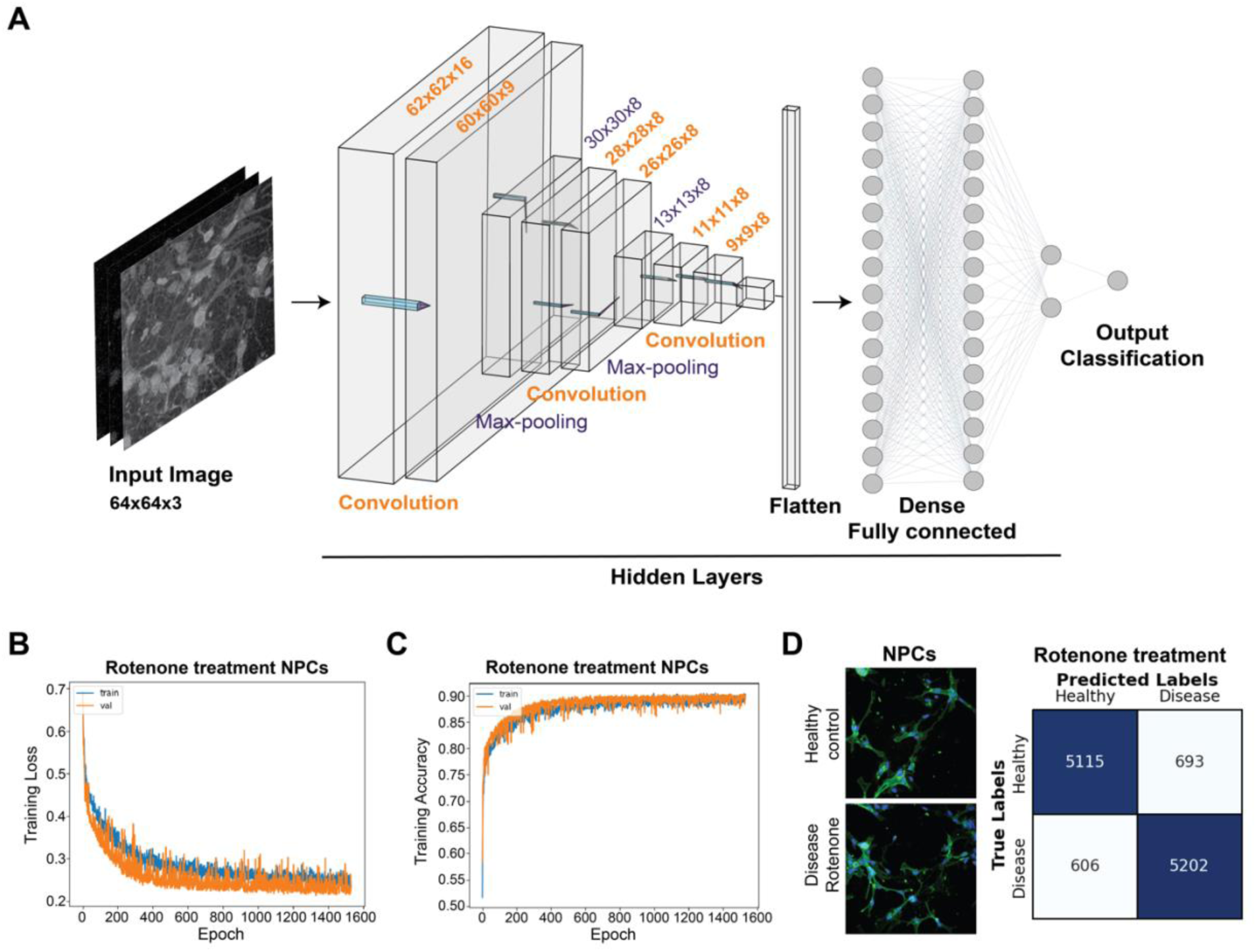
CNN models can be trained to predict PD disease status in a rotenone model of PD. **A)** Schematic of the binary CNN model, a simplified version of the VGG-16 model with 8 layers and 6042 trainable parameters, flattened to a layer of 16 nodes fully connected to a second layer of 16 nodes. The dimensional output for each convolutional and max-pooling layer are indicated. NPCs were treated with 1μM rotenone or vehicle for 20 hours. **B,C**) Line plot showing the training loss (B) and the accuracy (C) across epochs of training for classification of NPCs untreated or treated with rotenone. The training data is split into training (blue line) and validation sets (orange line). The model is trained and tested on both training and validation sets before updating in the next epoch. **D)** Right, example images of AIW002-02 NPCs untreated or treated with rotenone. Left, confusion matrix showing the predictions of the hidden test data predicted using the CNN trained with NPC treated/untreated with rotenone.

After successfully classifying a chemically induced model of PD, we next sought to investigate if we could classify a genetic model of PD while applying a similar approach as for rotenone. Homozygous mutations in the *Parkin* gene cause a form of autosomal recessive early onset PD.(14) The Parkin protein plays a key role in mitochondrial health, which is associated with all forms of PD.(15) For these tests, we compared the control iPSC line XCL1 with an isogenic line in which the Parkin gene was deleted, XCL1-ParkinKO.(9,16) NPCs were differentiated from the XCL1 and XCL1-ParkinKO iPSC lines, stained and imaged as above. For imaging of plates half of each 96 well plate was seeded with XCL1 NPCs and the other half with XCL1-ParkinKO NPCs, ensuring both lines were grown on the same dish for direct comparison. We also added mitotracker-red which binds to chloromethyl groups in the mitochondrial membrane as a third organic dye. The data was split into training and test datasets (80/20) as above, then used to train a CNN model (**Figure 2A**). We find the CNN classifier can distinguish XCL1 and XCL1-ParkinKO NPCs with an accuracy of 86.77 % (**Figure 2B**). We repeated this experiment in four separate batches of NPCs for a total of five batches. To test the robustness of the CNN model, we performed cross validation, training the model five times with different random splits within the training set into training and validation sets in each of the batch (**Figure 2C**). We found that for each experimental batch of NPCs the CNN model has a high accuracy and low variability in cross validation tests.

**Figure 2:**
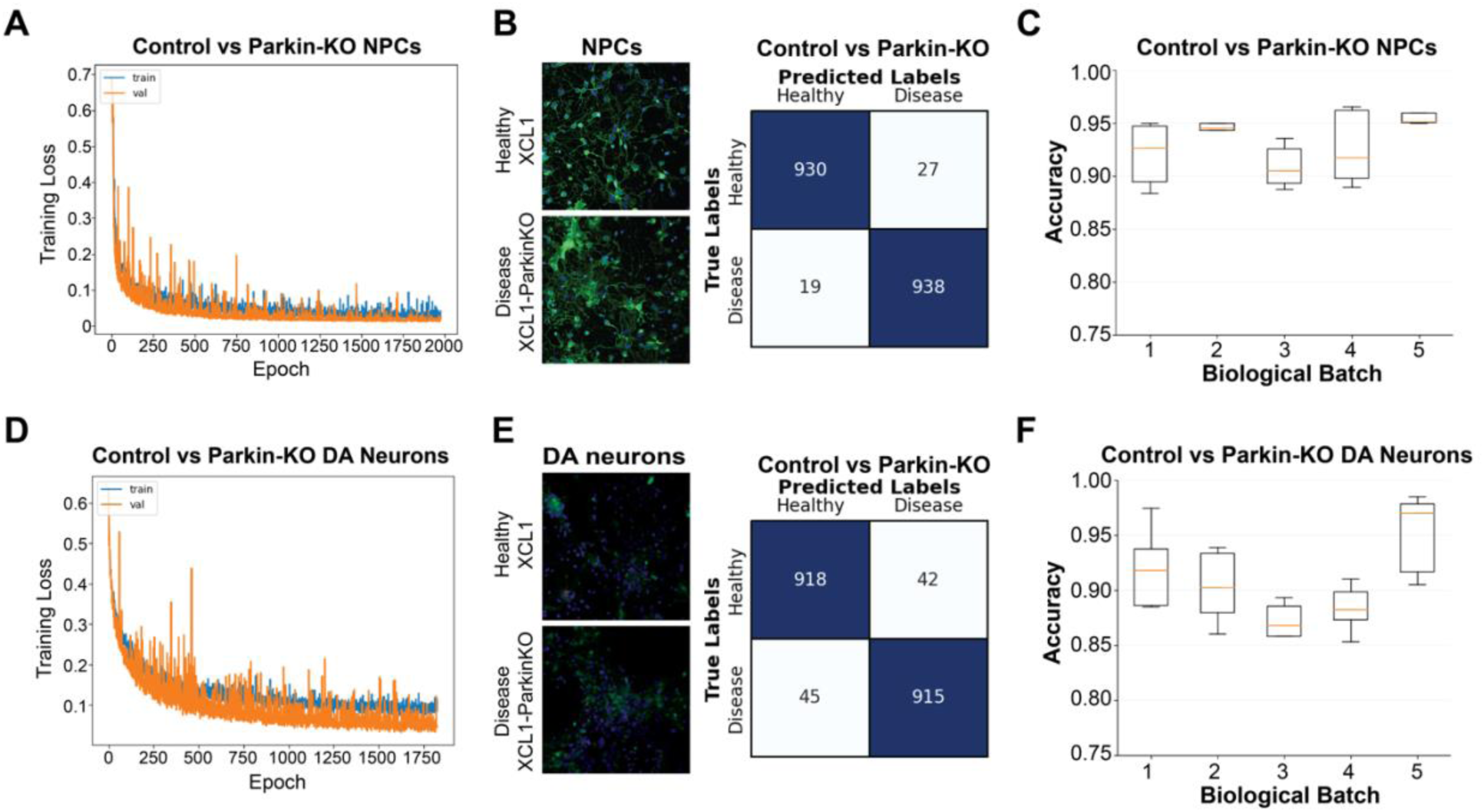
CNN models can be trained to predict PD disease status in a genetic model of PD, Parkin-KO compared to control. CNN models were trained using images of NPCs or DANs of control line XCL1 and PD model XCL1-Parkin-KO cells. **A,D)** Line plot showing the training loss across epochs of training a CNN to distinguish between Parkin-KO and control (A) NPCs and (D) DANs. **B,E)** Left, example images of XCL1 and XCL1-Parkin-KO (B) NPCs and (E) DANs. Right, confusion matrix of the prediction results for the text data in the CNN models trained to distinguish between control and Parkin-KO cells. **C,F)** Box plots showing the accuracy of CNN models trained on images of XCL1 vs XCL1-Parkin-KO (C) NPCs and (F) DANs with different random splits of training and validation sets. All models were tested on the same test data. Models were trained on 5 different biological replicates indicated on the x-axis.

After demonstrating that our CNN model can accurately predict whether NPCs are from the control or PD iPSC lines, we next explored whether this was applicable when dopaminergic neurons (DANs) were used instead of NPCs. Images of the DAN cultures were used to train CNN models with the same staining, processing and data splitting conditions applied to NPCs (**Figure 2D**). The CNN model for DANs has a very high accuracy of 95.47% (**Figure 2E**). Cross validation and experimental repeats show the CNN models are reproducible and robust for DANs (**Figure 2F**). Across both experimental repeats and cross validation with random start repeats we find the average accuracy for DANs models is 93.0% -/+ 2% and for DANs is 92.0% -/+ 5%.

### Combining nuclear, cell membrane and mitochondrial staining improves the accuracy of CNN models

In the Parkin KO and control CNN models, we utilized three input channels during classifier training. Nevertheless, the specific cellular components are driving the classifications is unknown when the models are trained with all three input channels. With this in mind, we set out to examine the individual contributions of the image information in each channel on the accuracy of our CNN classifiers. Hoechst (nuclei), GFP-WGA (cell membranes) and mitotracker-red (mitochondria) were imaged separately in different channels for both NPCs and DANs (**Figure 3A and B**). To assess the impact of each channel on the CNN model, we performed a series of experiments training CNNs with each channel in isolation and with combinations of each pair of channels. Each CNN training experiment was performed for five experimental batches with cross validation. For NPC images, CNNs trained with each channel alone were highly variable, with individual model accuracies ranging from 52-99% (**Figure 3C**). However, combinations of mitotracker-red with either Hoechst or WGA led to a high accuracy and low variation between repeats. For DAN images, CNNs trained with nuclear staining alone showed good accuracy, whereas CNNs trained with cell membrane or mitochondria staining alone have poor performance (**Figure 3D**). The mitochondria channel may not perform well in isolation as other contextual information may be required. For DANs, mitochondria are important to the classification as adding the mitochondria channel to models trained with the Hoechst or WGA improves the mean CNN classification by 15%. For both NPCs and DANs all channels together have the highest accuracy (**Figure 3C and D**). These findings collectively suggest that providing more information to the CNN enhances the classification accuracy.

**Figure 3:**
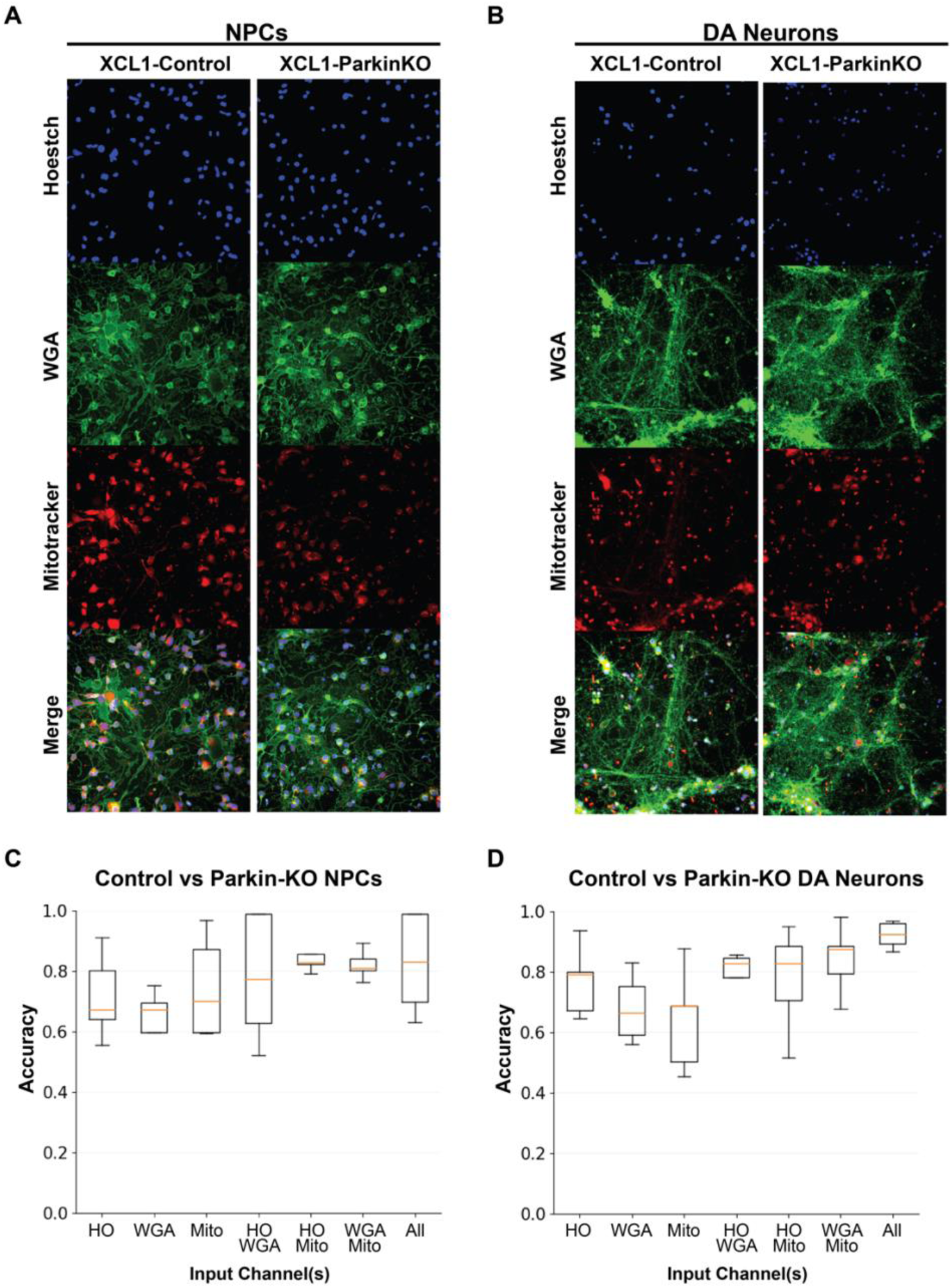
Nuclei, cell membrane and mitochondrial channels all contribute to CNN model accuracy. **A,B)** Example images of XCL1 control and XCL1-Parkin-KO (A) NPCs and (B) DANs with each channel shown separately and the merged images with all channels. Each channel has a different organic dye; Hoechst to make nuclei, WGA to stain the cell membrane and mitotracker-red for the mitochondria. **C,D)** Box plots showing accuracy of predictions of hidden test data using differently trained CNN models. CNN models were trained with each channel individually: Hoechst (HO), WGA, or mitotracker-red (Mito), or in pairs of two channels, or with all three channels together, indicated on the x axis. For each model the cross validation was performed on the training data. The experiment was performed on (C) NPCs and (D) DANs.

### Three different microscopy image processing methods produce highly accurate models

We next wanted to examine the effect of different image preprocessing methods on CNN classifier accuracy. In the models trained so far, images were processed by cropping and resizing the image to 64×64, and we refer to this method as “down sample”. To assess whether showing the CNN only the nucleus and cell soma would be sufficient to produce an accurate classifier, we used cell segmentation to detect nuclei and cropped a square around the radius of the nucleus, termed “cell segment” (**Figure 4A**). We trained a CNN using images processed by the “cell segment” method using the same number of input images as with the “down sample” processing method. For both NPC and DANs, the “cell segment” models were highly accurate (**Figure 4B and C**). The accuracy for hidden test data using the “cell segment” method for NPC images is 90.83%. For the DAN images the accuracy of the “cell segment” method is 94.42%. These finding indicate that machine vision can distinguish between PD and control cells using only the nucleus and cell soma.

**Figure 4:**
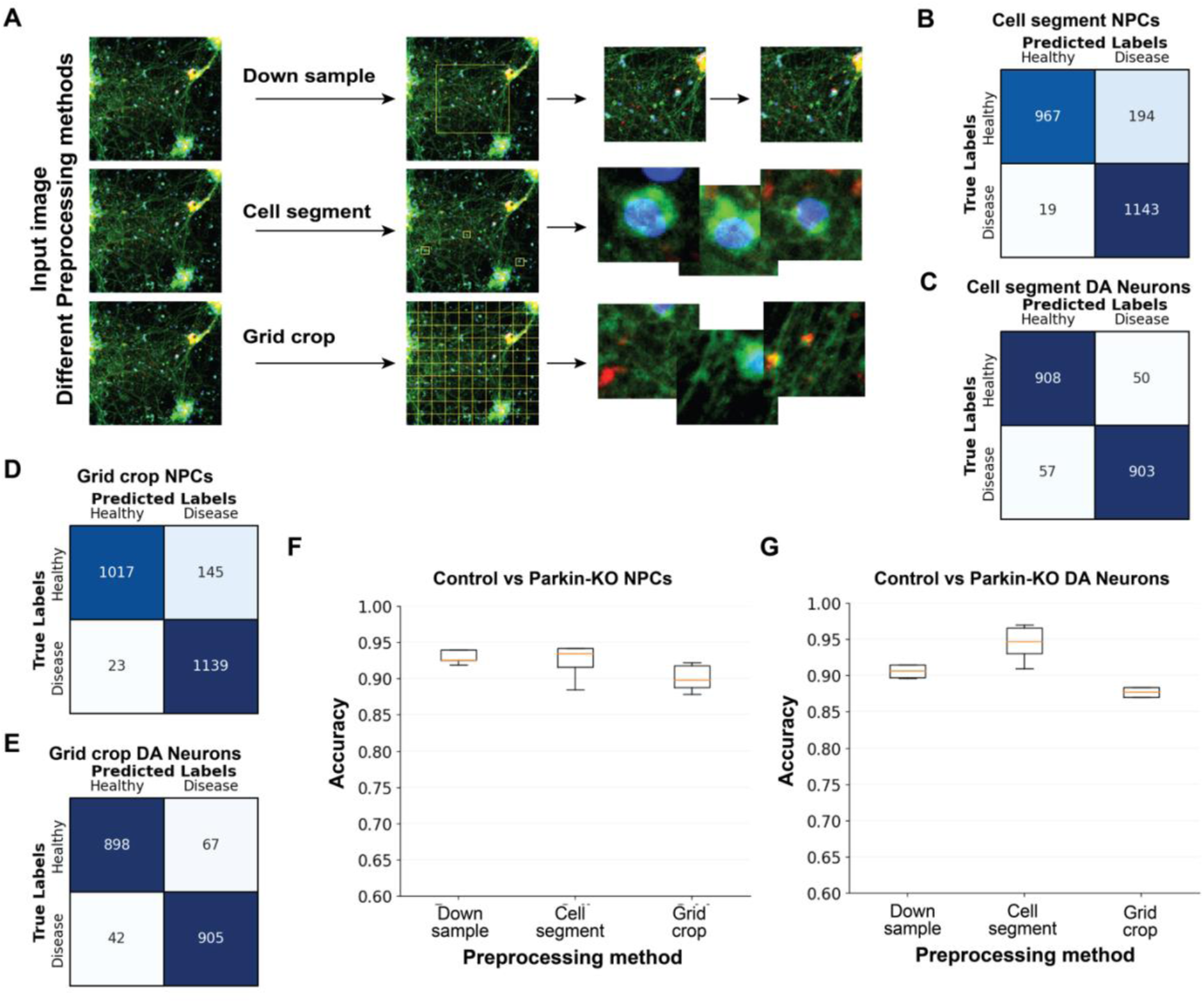
Input training data processed with different methods all yield accurate predictive models. **A)** Diagram with an example image processed with three different methods. The “down sample” method is the standard method used throughout the manuscript. The center of each image is cropped and then the resolution is reduced to result in a 64×64 pixel image. The “cell segment” method finds each nucleus in an image and crops an image sized relative to the nucleus. In the “grid crop” method the full image is split into 64×64 pixel images, in this method empty images were discarded. **B)** Confusion matrix showing the predicted disease status of test data with a CNN model trained to distinguish XCL1 to XCL1-Parkin-KO NPCs processed using “cell segmentation”. **C)** Confusion matrix showing the predictions of disease status images using a CNN model trained to distinguish XCL1 to XCL1-Parkin-KO DAN images processed using “cell segmentation”. **D)** Confusion matrix showing the predictions of test data with a CNN model trained to distinguish XCL1 to XCL1-Parkin-KO NPC images processed using “grid crop”. **E)** Confusion matrix showing the predictions of disease status in test data using CNN model trained to distinguish XCL1 to XCL1-Parkin-KO DAN images using “grid crop” processing. **F, G)** Box plot showing accuracy of test data predictions in cross validation experiments of different CNN models trained with images processed by three different methods, indicated on the x-axis. The experiment was performed on (F) NPCs and (G) DANs.

To test if distance between cells and cell processes could also contribute to the model, we tested cropping images in a simple grid (**Figure 4A**). We divided each starting image into 64×64 pixel images using a grid, that we refer to as “grid crop”. The resulting images have a wide variety of content. Some contain nuclei and cell bodies whereas others contain only neurites. Some images were empty, and we filtered these out by applying an intensity threshold. The “grid crop” images were used to train CNN models, including the same number of images as in the other methods. Again, both NPC and DAN models were highly accurate, 92.77% for NPCs and 94.30% for DANs using the “grid crop” method (**Figure 4D and E**). For the NPC images the “down sample” method yields the most accurate classifiers, indicating that including nuclear, cell soma, and spaces between cells are all required for the best accuracy (**Figure 4F**). In contrast, for DANs, the “cell segment” method produces CNN classifiers with the highest accuracy, possibly because the highly variable neurite processes are removed (**Figure 4G**). Overall, each of the three preprocessing methods result in models with accuracies over 90%.

### CNN models generalize within experiments but not across experiments

To test the ability of the CNN models to generalize, we trained five separate CNNs, one for each experimental batch of cells. We then tested each of the five separately trained models using the different hidden test image sets **(Figure 5A**). We find that for NPC images, each separately trained model is highly accurate for the test set from the same batch (as shown in Figure 1G). However, the disease status is poorly predicted in test images from other batches (**Figure 5B**). While in a few cases one model can predict if NPC images are of XCL1 or XCL1-Parkin-KO cells with an accuracy of 65%, in other cases the accuracies are equivalent or worse than random. For DANs, the ability of models trained with one experimental batch to predict if cells are from XCL1 or XCL1-Parkin-KO in a different batch is also poor (**Figure 5C**). We next tested if it was possible with this type of microscopy data for a model to generalize at all. For this we used four separate plates from one experimental batch, where all the plates were seeded, fixed, stained and imaged together. We trained a CNN for each plate and predicted if test images were from XCL1 or XCL1-Parkin-KO lines. For NPCs, all models predict the test data sets from all plates with high accuracy (**Figure 5D**), similar results were observed for DANs (**Figure 5E**). These results imply that the technical differences between experimental batches of NPCs and DANs are leading to CNN models that do not generalize. Indeed, within one experimental batch, CNN models generalize extremely well.

**Figure 5:**
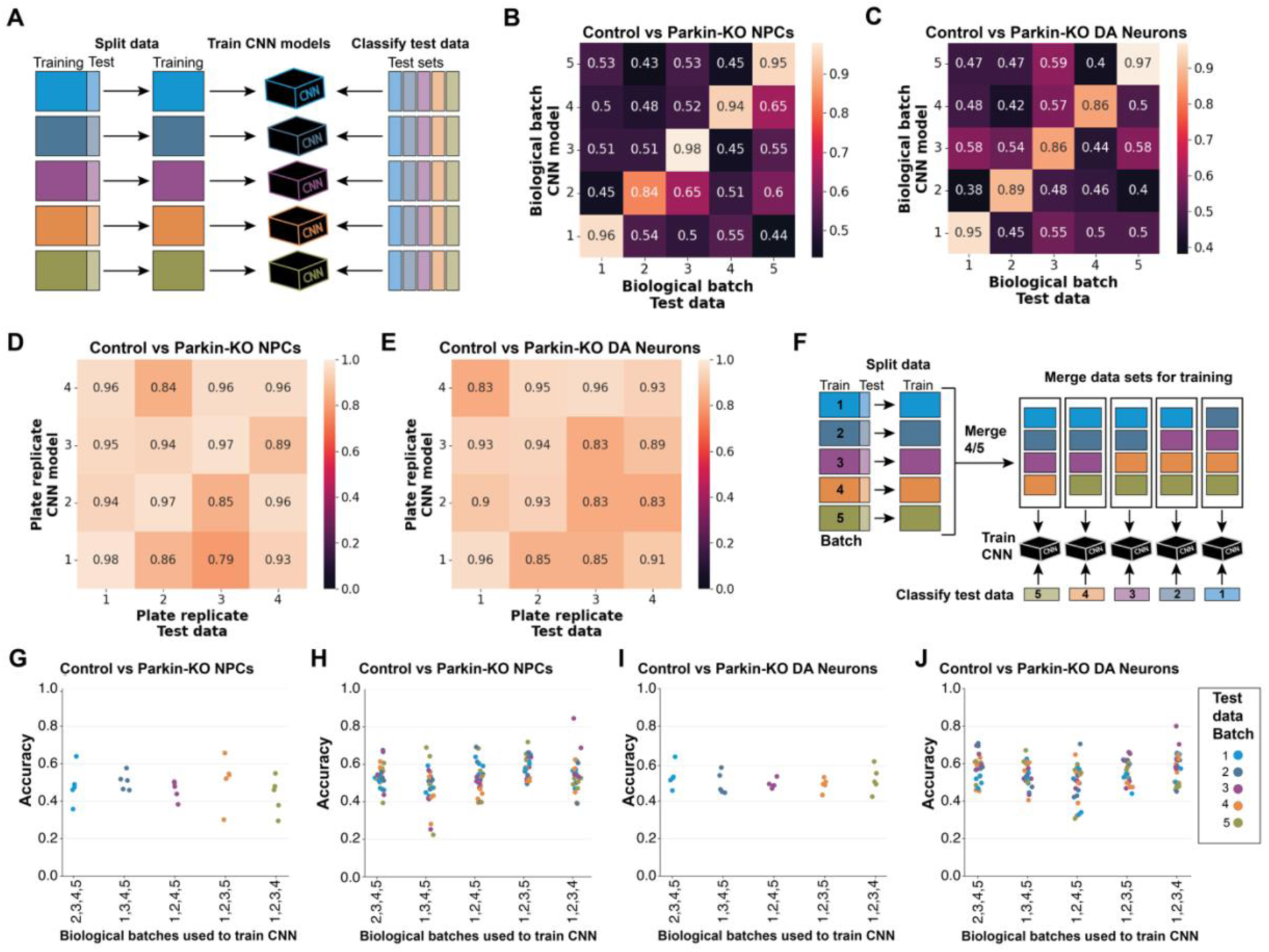
CNN models generalize within one batch of images but show poor generalization across batches. **A)** Schematic showing CNN models were trained separately for each of 5 batches (cells seeded, differentiated, stained, and imaged on different days). The test data for each of the 5 batches was predicted in all 5 separately trained model. **B, C)** Heat map showing the accuracy of prediction if images contain XCL1 control or XCL1-Parkin-KO (B) NPCs and (C) DANs. The y-axis indicates which batch was used to train the CNN model. The x-axis indicates which batch the predicted test data came from. **D, E)** Heat map showing the accuracy of prediction if images contain XCL1 control or XCL1-Parkin-KO (D) NPCs and (E) DANs. The y-axis indicates which plates was used to train the CNN model. The x-axis indicates the plate the predicted test data came from. **F)** Schematic showing batch drop out combinations, where a CNN model was training with 4 out of 5 batches and the test data is from the “dropped” batch. **G-H)** Dot plots showing the accuracy of CNN models trained with 4 of 5 batches. The test data used to train the models is shown on the x-axis. The batch the test data comes from is indicated by the colour of the dot. Each dot represents the same test data predicted in separately trained models; these replications represent different random splits within the training data. **G)** Accuracy of predictions of control vs Parkin-KO cells in NPC test images in the “dropped” batch not used in training. **H)** Prediction accuracy on all NPC test data sets from batch used in the training and the dropped batch for each CNN model. **I)** Accuracy of predictions of control vs Parkin-KO cells in DAN test images in the “dropped” batch not used in training. **J)** Prediction accuracy on all DAN test data sets from batch used in the training and the dropped batch for each CNN model.

With this in mind, we next set out to determine if combining multiple experimental batches and training CNNs with these merged data sets could help yield a model to accurately predict images from a data set not used in training (**Figure 5F**). We trained CNN models using NPC images from each combination of the four experimental batches and tested images from the dropped-out batch, for each combination we repeated the training five times (**Figure 5G and H**). Next, we performed the same experiment on DANs (**Figure 5I and J**). We find that for both NPCs and DANs the models occasionally have an accuracy near 70% in the test data from the dropped batch, but more often have accuracies near or below 50%. Given that some models tend toward generalization, we decided to test tuning different hyperparameters, step size and learning rate. We found in the ranges we tested that altering either step size or learning rate did not improve the ability of a CNN trained on four experimental batches to generalize to the fifth batch (data not shown).

### CNN models can accurately distinguish between isogenic controls and different genetic models of PD

We have shown that a CNN can accurately predict if images are of NPCs or DANs from control or Parkin KO iPSCs in one genetic background, the XCL1 cell background. Thus, we next wanted to confirm that the distinction is not iPSC line specific. Building on the earlier work, we used a second healthy control line (referred to as AIW002-02) and compared cells generated from this control line to cells generated from this line with Parkin knocked out.(17) The iPSCs were differentiated into NPCs, stained, and imaged as described above. Here we have ∼3600 images per condition. A CNN model was trained with AIW002-02 and AIW002-02-ParkinKO NPCs, and we find the prediction accuracy is 95.34% on the hidden test data (**Figure 6A**). After confirming Parkin-KO cells can be distinguished from control cells across two different genetic backgrounds, we next tested two other genetic models of PD. The gene *GBA1* encodes the enzyme GCase, critical for lysosomal function and cellular homeostasis. Mutations in *GBA1* constitute a major risk factor for PD.(18) We trained a CNN model to distinguish between AIW002-02 control NPCs and AIW002-02-GBA1-KO NPCs. The model predicts the correct iPSC line with 86.58% accuracy (**Figure 6B**). Mutations or triplication of the gene SNCA encoding α-synuclein is a common genetic cause of PD.(14) We used a line with the known patient mutation A53T introduced into the SNCA gene in AIW002-02 iPSC line, AIW002-02-SNCA-A53T. When we trained the CNN model with NPC images from AIW002-02 and AIW002-02-SNCA-A53T we find the model can distinguish the two cell lines with 97.81% accuracy on the hidden test data (**Figure 6C**).

**Figure 6:**
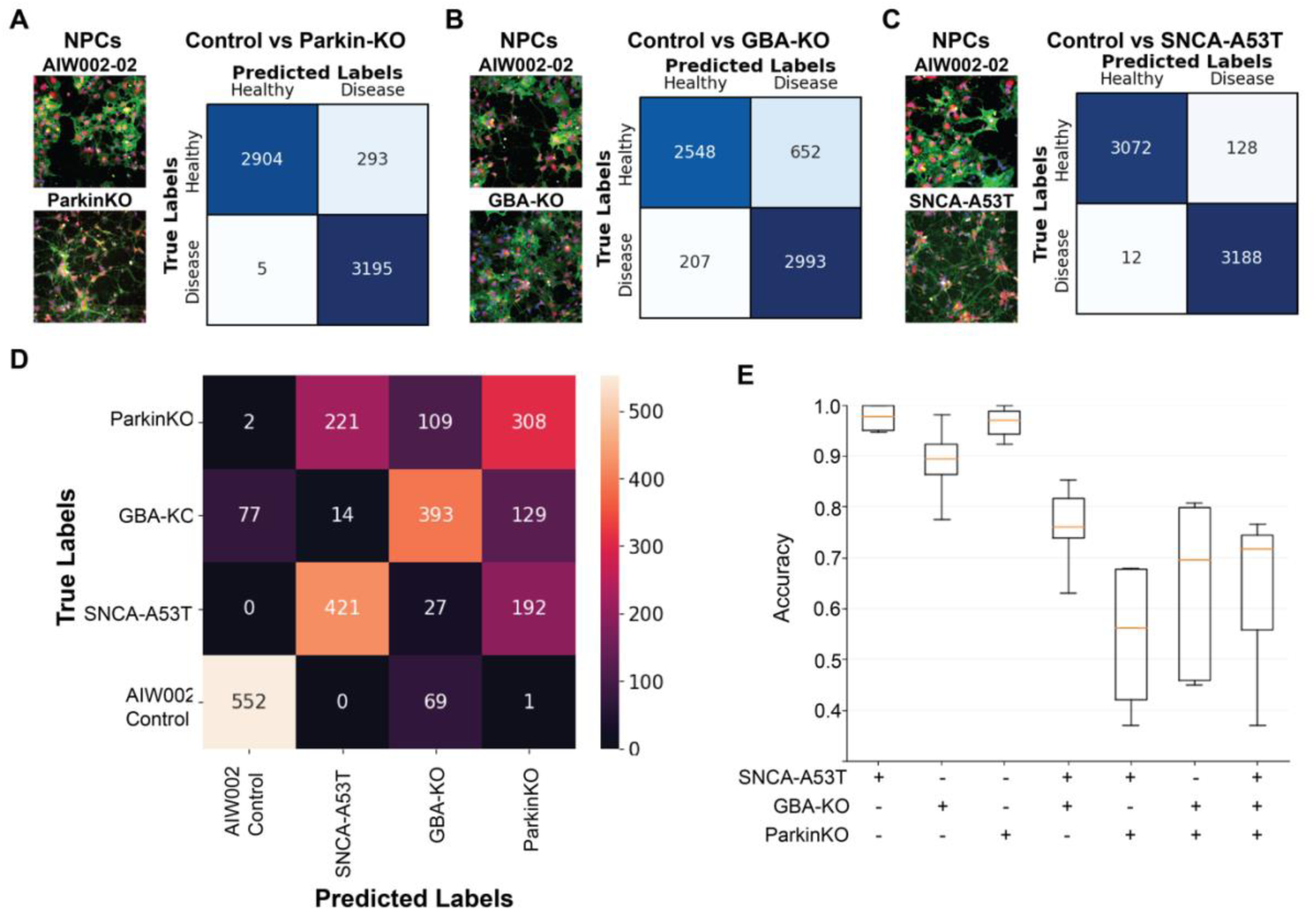
CNN classifiers accurate distinguish between control NPCs and three different models of PD. **A)** Right, example images of control AIW002-02 and AIW002-02-Parkin-KO NPCs. Left, confusion matrix showing predictions of hidden test data in a model trained to distinguish AIW002-02 (healthy) and AIW002-02-Parkin-KO (disease) NPCs. The number of images predicted are indicated. **B)** Right, example images of control AIW002-02 (healthy) and AIW002-02-GBA-KO (disease) NPCs. Left, confusion matrix showing the number of images from the hidden test data predicted as healthy or control and the true status of the cell. **C)** Right, example images of control AIW002-02 (healthy) and AIW002-02-SNCA-A53T (disease) NPCs. Left, confusion matrix showing the number of images from the hidden test data predicted as healthy or control and the true status of the cell. **D)** Confusion matrix showing test data predictions of which cell type an NPC images contains. A four-way categorical model was trained to distinguish between AIW002- 02 control, AIW002-02-SNCA-A53T, AIW002-02-GBA-KO and AIW002-02-ParkinKO NPCs. **E)** Box plot showing accuracy of cross validation of CNN binary classier models trained with different combinations of inputs. The x-axis indicates which disease model NPC images were compared to AIW002-02 control cells. Each line AIW002-02-SNCA-A53T, AIW002-02-GBA-KO and AIW002-02-ParkinKO was used to train CNN models alone, in combination of two PD lines together and all three PD lines together. For each combination of iPSC lines cross validation replication with random splits of the training data was performed five times.

To explore the extent of overlap in morphology between the three genetic models of PD we designed a four-way categorical model to classify between control, Parkin-KO, GBA-KO and SCNA-A53T NPCs. The architecture of the CNN was maintained, only the final output layer was adjusted to account for four output categories instead of two. We trained the categorical model with NPC images from AIW002-02, AIW002-02-Parkin-KO, AIW002-02-GBA-KO and AIW002-02-SNCA-A53T, then tested the accuracy of predictions in the hidden test data. Each image is predicted as being from one of the four iPSC lines. We find that overall accuracy is only 66.56%, however the accuracy for predicting the AIW002-02 control line correctly is 88.75% (**Figure 6D**). The categorical CNN model mis-predicts the different genetic PD models as another PD model but does not confuse them for the control line. This indicates a partially overlapping PD morphology in these lines. Specifically, AIW002-02-GBA-KO and AIW002-02-SNCA-A53T are both mis-predicted as AIW002-02-ParkinKO, but not mis-predicted as each other. In the cases where AIW002-02 is predicted as a PD line, it is predicted to be the AIW002- 02-GBA-KO, indicating this line might share the most morphological similarity to the control.

After noting that the different PD genetic models could have overlapping morphological features used by the CNN, we decided to test combinations of the three genetic PD models in training CNN binary classifiers. Surprisingly, when images from AIW002-02-GBA-KO and AIW002-02-SNCA-A53T were combined together to train a CNN model to distinguish these two genotypes from the AIW002-02 control, the mean accuracy of predicting the hidden text data is 76.5%, indicating these two models of PD may have overlapping morphological phenotypes recognized by the CNN (**Figure 6E**). However, the models trained by combining sets including AIW002-02-Parkin-KO together with AIW002-02-GBA-KO or AIW002-02-SNCA-A53T or both together are highly variable over cross validation, indicating the variability between genotypes and within one genotype is very high **(Figure 6E**). Our findings indicate overlap between morphological difference in multiple genetic models. CNN Models distinguishing individual PD genetic models from control cells are all highly accurate and perform better than models trained on two or more genetic PD models.

## Discussion

In this study, we introduced a deep learning convolutional neural networks (CNNs) to detect disease status in iPSC-derived cells, specifically NPCs and DANs. Notably, we harnessed the power of high-content imaging using three distinct organic dyes to stain the nucleus, cell membrane, and mitochondria. While previous research has employed CNNs in the context of microscopic imaging data, these applications were primarily focused on cell type classification or cell segmentation.(19,20) High content microscopy analysis methods targeted toward screening applications utilize extracted features and do not directly apply machine vision to the microscopy images.(21,22) While, to our knowledge, there exists one other notable instance of CNN application for disease status classification in Parkinson’s disease (PD), it involved patient fibroblasts rather than neurons.(23) In contrast, our study takes a significant step forward by utilizing CNNs to distinguish disease status in iPSC-derived DANs, thereby expanding the utility of this technology in the field of neurobiology.

In a related study, the Gandhi group utilized features extracted from microscopy images to train a deep learning neural network for predicting disease status in iPSC-derived cortical neuron cultures.(24) Their research, which employed an SNCA triplication line, provided valuable insights into disease classification. However, our approach, which incorporates CNNs and directly analyzes microscopy images, offers a distinct contribution to this evolving field. Notably, our study demonstrates that CNN models can be effectively trained to predict disease status, even in complex biological systems.

To validate the effectiveness of our CNN classifier, we employed two distinct models of PD: a pharmacological model involving rotenone treatment and a genetic model using several lines with PD-associated mutations or gene knockouts. Importantly, our approach successfully distinguished disease status in isogenic control and Parkin KOs in both NPCs and DANs, providing evidence of its versatility and robustness. Moreover, we extended our analysis to NPCs and DANs differentiated from isogenic iPSC lines with mutations in *SNCA* and *GBA*, two additional PD genes, highlighting the potential of CNNs to classify disease status across various disease models.

To ensure the reproducibility of our model, we subjected it to rigorous testing, including cross-validation and validation over multiple biological datasets. These tests confirmed that our model is both computationally and biologically reproducible, underscoring its reliability and utility. Additionally, we investigated the contributions of different channels within the imaging data, revealing that all three channels, representing the nucleus, cell membrane, and mitochondria, play vital roles in the model’s accuracy, with the nucleus channel being the most important contributor to predictions in our model.

One limitation of CNNs is that they can classify based on irrelevant information. We explored the effects of image processing on model performance, comparing segmenting individual cells and dividing large images into a grid of smaller ones with the traditional method of cropping and resizing. Our findings suggest that CNNs rely on the information provided by the nucleus and cell body to make accurate predictions. This insight provides valuable information about the features that contribute most to disease status classification and indicates biologically relevant information is being used for classification.

One notable challenge we encountered was the generalizability of our model across different biological experiments. While some degree of generalization was observed, it was not consistently achieved. This lack of generalizability appeared to stem from the inherent variability between batches of NPCs and DANs, a well-known occurrence in biological research. Addressing this challenge may require further adjustments to the image processing method to normalize images and enhance the model’s capacity to generalize across varying culture conditions.

In summary, our CNN-based model presents a valuable tool for the exploration of nervous system biology when tested alongside a range of CNN models. The binary predictive models developed in this study have promising applications in phenotypic compound screening. For instance, researchers can employ CNN models to differentiate control cells from PD models and simultaneously treat PD cell models with a panel of drugs or compounds of interest. Such an approach allows for the identification of compounds that can cause PD-affected cells to be classified as the control state, indicating a potential rescue of the PD phenotype. Conversely, researchers can perform a reverse screen to identify compounds that lead to control cells being predicted to be more closely aligned with a PD state, shedding light on potential factors that drive a PD phenotype. These applications underscore the versatility and significance of our CNN-based disease status classifier in advancing our understanding of PD and contributing to potential therapeutic interventions.

## Methods

### Cell lines

The iPSC lines XCL1 and XCL1-ParkinKO were purchased from Xcell Science and previously described.(9) The XCL1 line was reprogrammed from cord blood cells from a male donor using episomal vectors. The other AIW002-02 and AIW002-02-Parkin-KO iPSC cell lines were previously described.(12,17) The AIW002-02-GBA-KO and AIW002-02-SNCA-A53T lines were generated using CRISPR and standard quality control measure were performed (see supplemental documents 1 and 2).

AIW002-02-SNCA-A53T: A single guide RNA (gRNA) was designed using benchling.com to generate a double-strand break (DSB) in exon 3 (ENSE00000970012) of transcript SNCA-201 (ENST00000336904.7) of the human gene *SNCA* (ENSG00000145335), 1 base pair (bp) downstream of the target nucleotide. The mutation A53T was created by homology-directed repair (HDR) using a single-stranded oligonucleotide (ssODN) template. All CRISPR reagents were electroporated into iPSCs using the P3 Primary Cell 4D Nucleofector™ X Kit S (Lonza), as previously described.(25) Cells at 50% confluency were dissociated with Accutase (StemCell Technologies). 500,000 cells were resuspended in 25 μl of Cas9:gRNA ribonucleoprotein (RNP)-ssODN-buffer mix, consisting of 1 µl of Alt-R® S.p. HiFi Cas9 Nuclease V3 (stock 61 µM; IDT), 3 µl of gRNA (stock 100 µM; Synthego), and 1 µl of ssODN (stock 100 µM; IDT) in 20 µl of nucleofection buffer P3. Nucleofection was performed using the CA137 program in a Nucleofector 4D device.

Following nucleofection, iPSCs were evenly distributed into a flat-bottom 96-well plate in mTeSR1 media (StemCell Technologies) and 10 µM Y-27632 (StemCell Technologies). After limiting dilution, gene-edited clones were identified by ddPCR (QX200™ Droplet Reader, Bio-Rad). The detection of the modified nucleotide by ddPCR was based on a TaqMan® assay including two PCR primers and two DNA probes (one for each of the wildtype and mutant alleles). Locked Nucleic Acid (LNA®) probes were designed following the manufacturer’s criteria. Sequence integrity of successful clones was assessed using Sanger sequencing of PCR amplification of the expected 320 bp amplicon (**Supplemental document 1**).

AIW002-02-GBA-KO: Two gRNAs were designed using the “optimized CRISPR design” tool (www.crisp.mit.edu) to target sites in exon 4 of the human *GBA* gene (ENSG00000177628) and remove a 98bp segment. Each double strained gRNA was separately cloned into a Cas9/puromycin expressing vector (pX459 from Addgene #48139). The gRNA plasmids were delivered into AIW002- 02 iPSCs by Neon electroporation. Cells recovered for 48hours and then 0.2µg/ml Puromycin was added into medium to select transfected cells and isolate clones. Cell clones were transferred in 96 well plates for genotyping and amplification.

KO cells were confirmed by PCR amplification and gel electrophoresis, where the wildtype GBA gene results in a 739bp product and the KO product was 641bp. Genomic DNA was extracted with QuickExtract (Lucigen) and PCR was performed using Q5® High-Fidelity DNA Polymerase according to the manufacturer’s protocol. PCR products were cloned into the pMiniT vector and sent for Sanger sequence to confirm the KO by the DNA sequence (**Supplemental document 2**).

**Table 1:**
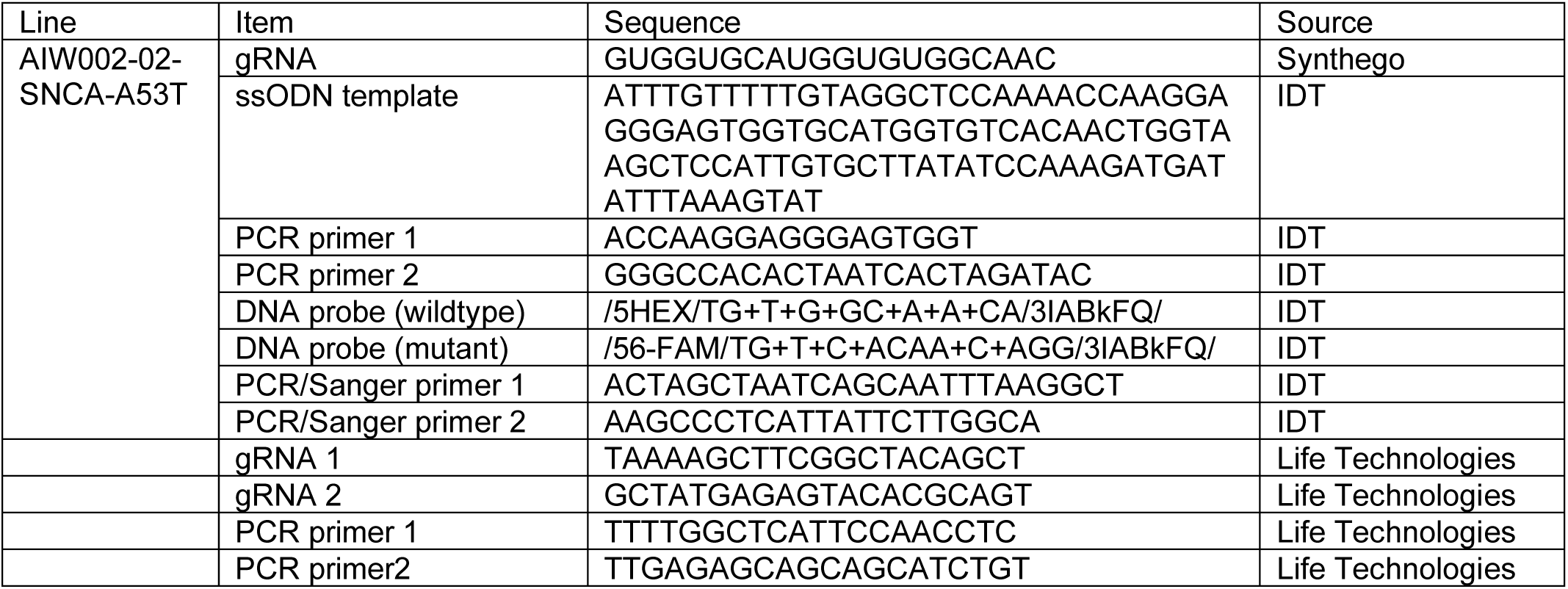
Sequences of reagents used in gene editing to generate new genetic PD model iPSC lines.

### Cell cultures and iPSC differentiation

IPSCs were differentiated into NPCs as previously described and frozen for future experiments.(12) NPCs were recovered for one week and seeded at a density of 10,000 cells per well in 96-well plates coated with poly-L-ornithine (PLO) 1μg/ml overnight at 37 ^0^C, followed by laminin 5μg/ml for 2 hours at 37 ^0^C. NPCs were stained and fixed 48 hours after seeding. DANs differentiated from NPCs as previously described and maintained in 96 well plates for 4 weeks.(17)

### Rotenone treatment

NPC were seeded at a density of 10 000 cells per well in 96-well plates. After 24 hours for cell adhesion and recovery, half of the wells were treated with 1uM rotenone for 20 hours. Rotenone treatment was selected from visual inspection of a rotenone dose curve and from previous experiments, to select a dose that would not cause a visual difference in cell to the human eye.(26)

### Fixation and staining

To stain for mitochondria, live NPCs and DANs were incubated with 500nM Mitotracker OrangeCMTMRos (ThermoFisher Scientific) for 1 hour at 37 ^0^C. Then the cells were fixed with 4% formaldehyde in PBS (10 mins), washed 3 times in PBS, stained with CF488A conjugated Wheat germ agglutinin (Biotium, 2μg/mL) and Hoechst33342 (ThermoFisher Scientific, 5μg/mL) for 10 minutes and washed 3 times in PBS.

### Imaging

Images were acquired on a CellInsight CX7 High Content Screening microscope (ThermoFisher Scientific) with a 20X objective (NA 0.7). Excitation/ Emission filters for Hoechst33324, WGA and Mitotracker were 386-23/438-47, 485-20/542-27, 549-15/612-69, respectively. Images were collected for 121 fields per well (covering ∼90% of a given well). Image size was 1104×1104 pixel (0.4µm/pixel) with 14-bit depth. For analysis, images were exported in 8-bit TIFF format.

**Table 2:** Sets of microscopy images from cells seeded, grown, fixed, and imaged at different times.

| **Experiment** | **Batch** | **iPSC lines** | **Cell type** | **Number of 96 well plates** | **Cell staining** |
| --- | --- | --- | --- | --- | --- |
| -/+ 1 mM Rotenone | 1 | XCL1 | NPCs | 6 | Hoechst, WGA |
| Control vs Parkin-KO | 1 | XCL1, XCL1-Parkin-KO | NPCs | 2 | Hoechst, WGA, mitotracker |
| Control vs Parkin-KO | 2 | XCL1, XCL1-Parkin-KO | NPCs | 2 | Hoechst, WGA, mitotracker |
| Control vs Parkin-KO | 3 | XCL1, XCL1-Parkin-KO | NPCs | 4 | Hoechst, WGA, mitotracker |
| Control vs Parkin-KO | 4 | XCL1, XCL1-Parkin-KO | NPCs | 4 | Hoechst, WGA, mitotracker |
| Control vs Parkin-KO | 5 | XCL1, XCL1-Parkin-KO | NPCs | 2 | Hoechst, WGA, mitotracker |
| Control vs Parkin-KO | 1 | XCL1, XCL1-Parkin-KO | DANs | 2 | Hoechst, WGA, mitotracker |
| Control vs Parkin-KO | 2 | XCL1, XCL1-Parkin-KO | DANs | 2 | Hoechst, WGA, mitotracker |
| Control vs Parkin-KO | 3 | XCL1, XCL1-Parkin-KO | DANs | 4 | Hoechst, WGA, mitotracker |
| Control vs Parkin-KO | 4 | XCL1, XCL1-Parkin-KO | DANs | 4 | Hoechst, WGA, mitotracker |
| Control vs Parkin-KO | 5 | XCL1, XCL1-Parkin-KO | DANs | 2 | Hoechst, WGA, mitotracker |
| Control vs Parkin-KO | 1 | AIW002-02, AIW002-02-Parkin-KO | NPCs | 18 wells per condition | Hoechst, WGA, mitotracker |
| Control vs GBA-KO | 1 | AIW002-02, AIW002-02-GBA-KO | NPCs | 18 wells per condition | Hoechst, WGA, mitotracker |
| Control vs SNCA-A53T | 1 | AIW002-02, AIW002-02-SNCA-A53T | NPCs | 18 wells per condition | Hoechst, WGA, mitotracker |

For rotenone experiments on each plate half the wells were treated with 1 mM Rotenone and half with vehicle. For XCL1 vs XCL1-Parkin-KO experiments on each plate half the wells were seeded with XCL1 and half with XCL1-Parkin-KO. For AIW002-02 experiments three columns were seeded for each genotype (AIW002-02, AIW002-02-Parkin-KO, AIW002-02-GBA-KO and AIW002-02-SNCA-A53T) in a 96 well plate.

### Image preprocessing

Down sample method: The basic preprocessing methods the input images 1104 by 1104 pixels were down sampled to a 128×128 pixel image with linear interpolation, and then cropped in the margins by another 32 pixels to get an input image of 64×64 pixels. Each channel is process separately. This method was applied for all models unless otherwise indicated.

Cell segment method: the python package PIL was used to identify nuclei and measure the diameter of the nucleus. Then a 64 x 64 box is drawn around the nucleus with the center of the nucleus set as the center of the box.

Grid crop method: The original images of 1104×1104 were cropped into 144 images and rescaled with linear interpolation to 64×64 pixel images. Intensity was used to detect empty images after cropping. For all images, preliminary preprocessing was used to find and reject images that were empty or contained the edge of the plate. A brightness threshold, the top and bottom 10% average pixel intensities was used remove images and reduce variation in the data set. All other images had each of their channels normalized, before entering the CNN model.

### CNN model

We used the VGG16 changing from 16 layers to 8 layers. The model was created in Python using the Keras deep learning library running on the TensorFlow framework. An 8-layer convolutional model was created, incorporating 3 blocks of convolutional and pooling layers with the rectified linear unit (ReLU) activation function followed by a fully connected classification perceptron with the SoftMax activation function.

For the categorical model, the overall structure of the model remains the same, however, changes were made to the final SoftMax activation layer to support multiple labels with categorical cross entropy.

### CNN training

Processed images were split 80/20 into training and test images. The training set was then split again 80/20 into training and validation. The CNN default model conditions are set as 64 steps per epoch for 50 epochs. Separate models were created to handle different amount of channel inputs, differing only in the first convolutional layer to handle the different number of channels. The experiments involving testing different channel combinations used models for each of the indicated channels. For the rotenone treatment models only two channels (nuclei and cell membrane) were imaged and used as inputs to the CNN. For all other experiments three channels were used.

**Table 3:** Number of images used in training different models before data splitting.

| CNN model | Number of images used in each condition |
| --- | --- |
| AIW002-02 Rotenone vs control NPCs | 34848 |
| XCL1 control vs XCL1-Parkin-KO NPCs | 5808 |
| XCL1 control vs XCL1-Parkin-KO DANs | 5808 |
| AIW002-02 control vs AIW002-Parkin-KO | 3872 |
| AIW002-02 control vs AIW002-Parkin-KO | 3872 |
| AIW002-02 control vs AIW002-Parkin-KO | 3872 |

### Cross validation

For cross validation experiments within a given dataset, we divided the training dataset into 5 equal-sized subsets. The model is then trained on 4 of the subsets and evaluated on the remaining subset, which serves as the validation set. This process is repeated 5 times, with each subset serving as the validation set once. The performance of the model is then averaged across the 5 iterations to obtain a more robust estimate of its generalization performance.

### Dropout comparisons for generalization

For experiments where four batches were used as the training data and the dropped batch was used as the test data. The same processed images used in separate batch models were used for the generalization tests.

### CNN model outputs

For each model training curves for error loss and accuracy are output. Average receiver operating characteristic (ROC) curves and accuracy are calculated from the test results. Outputs are printed to a log file and plots are saved. Accuracy is a calculated by the amount correctly classified labels over the total amount of items.

### Code availability

Cod for image processing, building models, training models and testing models is freely available at: https://github.com/RhalenaThomas/DeepLearningCNN_DiseaseStatusClassifier

### Author Contributions

Concept by RAT and EAF. Cell cultures by EM, CXQC, CH and RL. Cell staining and high content imaging by AK and WR. Generation of CRISPR iPSC lines by ED and ZY. CNN model architecture, coding and running experiments by EC. Preprocessing methods and model testing by SS. Experimental design by EC and RAT. Data interpretation by RAT and EC. The study was supervised by TMD and EAF. The manuscript was written by RAT, all authors reviewed, edited, and agreed on the contents.

## Acknowledgments

This work was supported a CIHR Foundation grant (FDN – 154301), a Fonds d’Accéleration des Collaborations en Santé (FACS) grant from CQDM/MEI awarded to EAF and a project grant from CIHR (PJT-169095) awarded to TMD. EAF is a Canada Research Chair (Tier 1) in Parkinson’s disease. TMD received funding through the McGill Healthy Brains for Healthy Lives (HBHL) initiative, the CQDM Quantum Leaps program with support from Brain Canada, the Alain and Sandra Bouchard Foundation, the Sebastien and Ghislaine Van Berkom Foundation, Médicament Québec, and the Mowafaghian Foundation. RAT received funding through the McGill Healthy Brains for Healthy Lives (HBHL) Postdoctoral Fellowship and Molson NeuroEngineering Fellowship.

---

**Supplemental Document 1.**
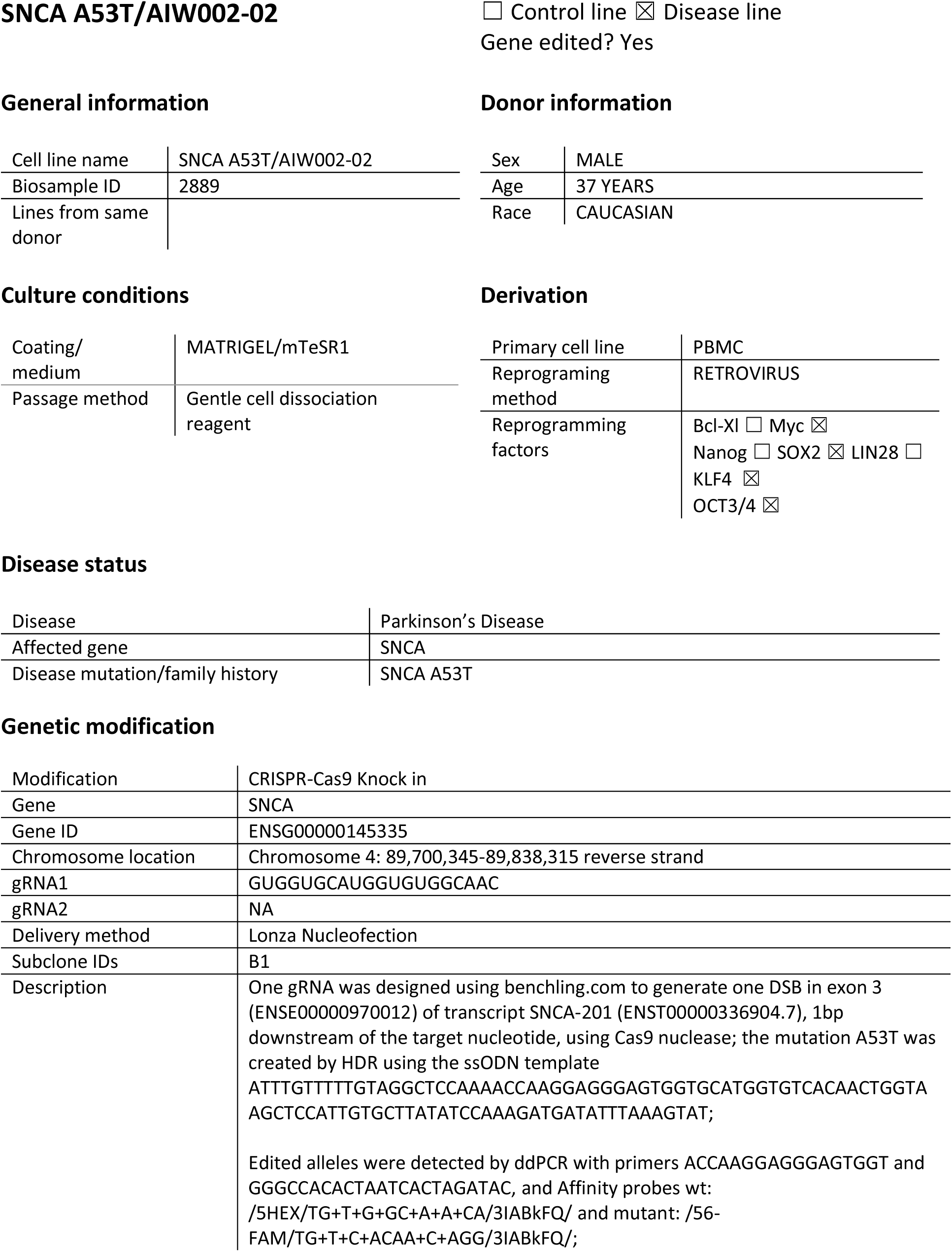
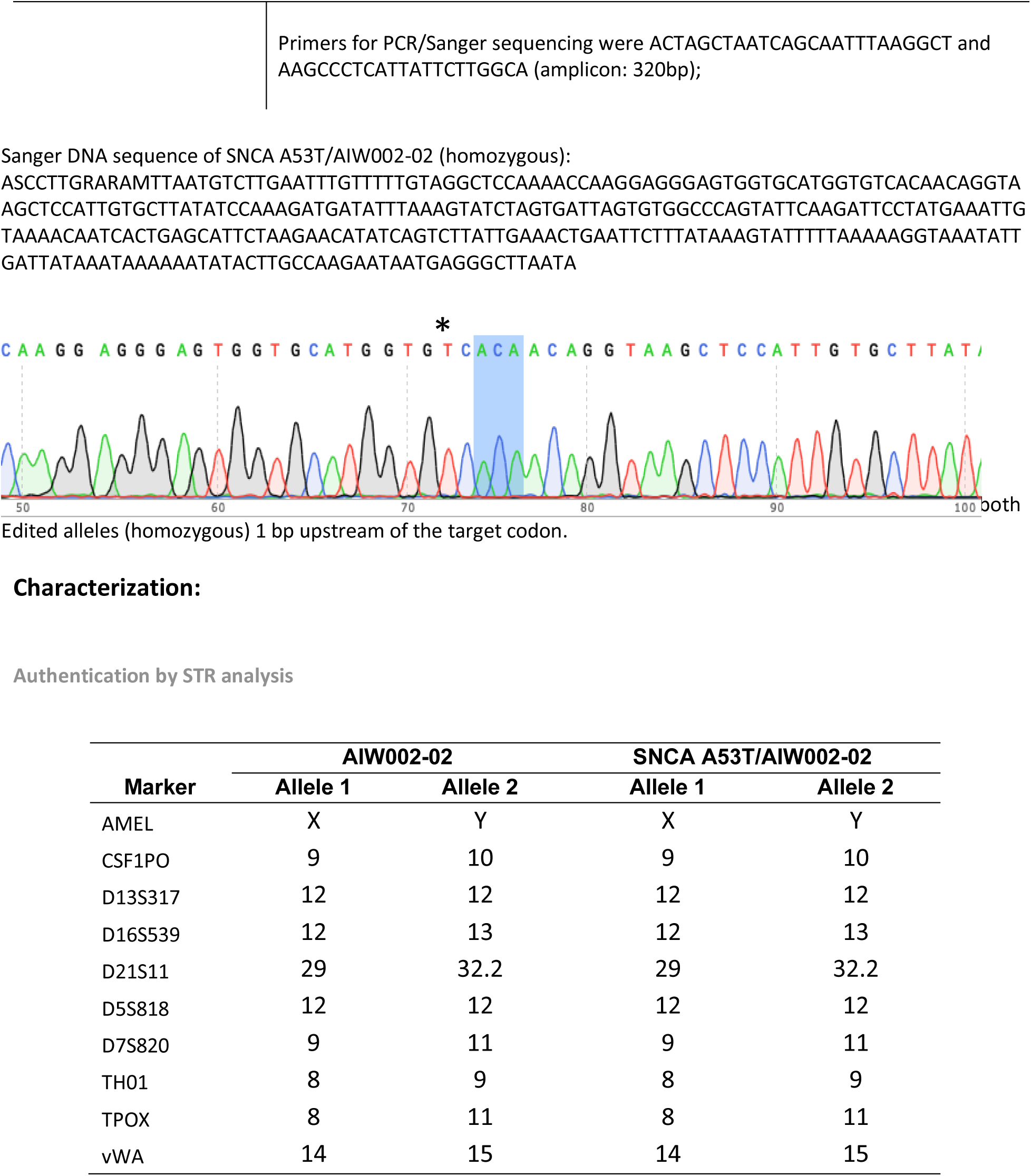
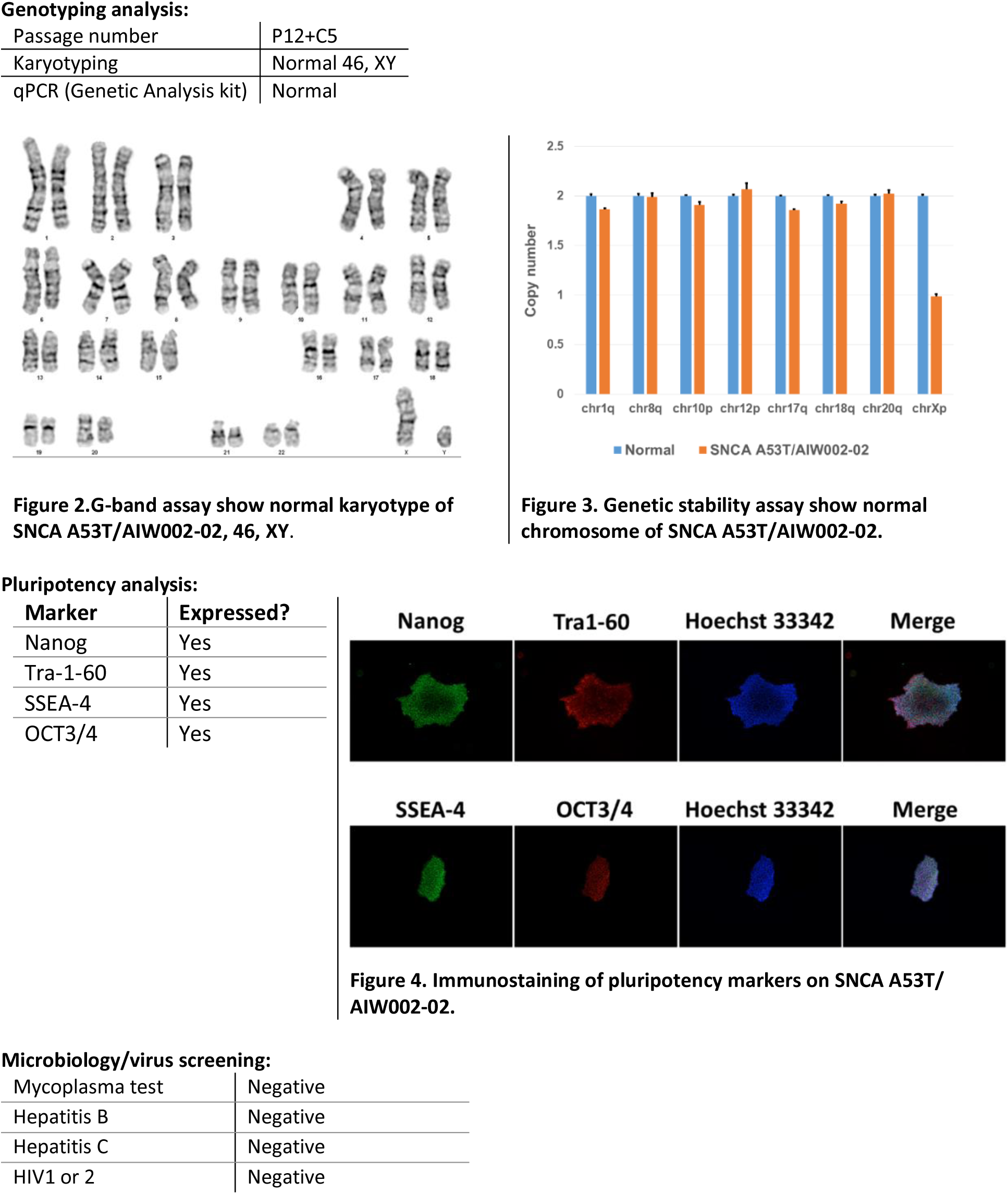

**Supplemental Document 2.**
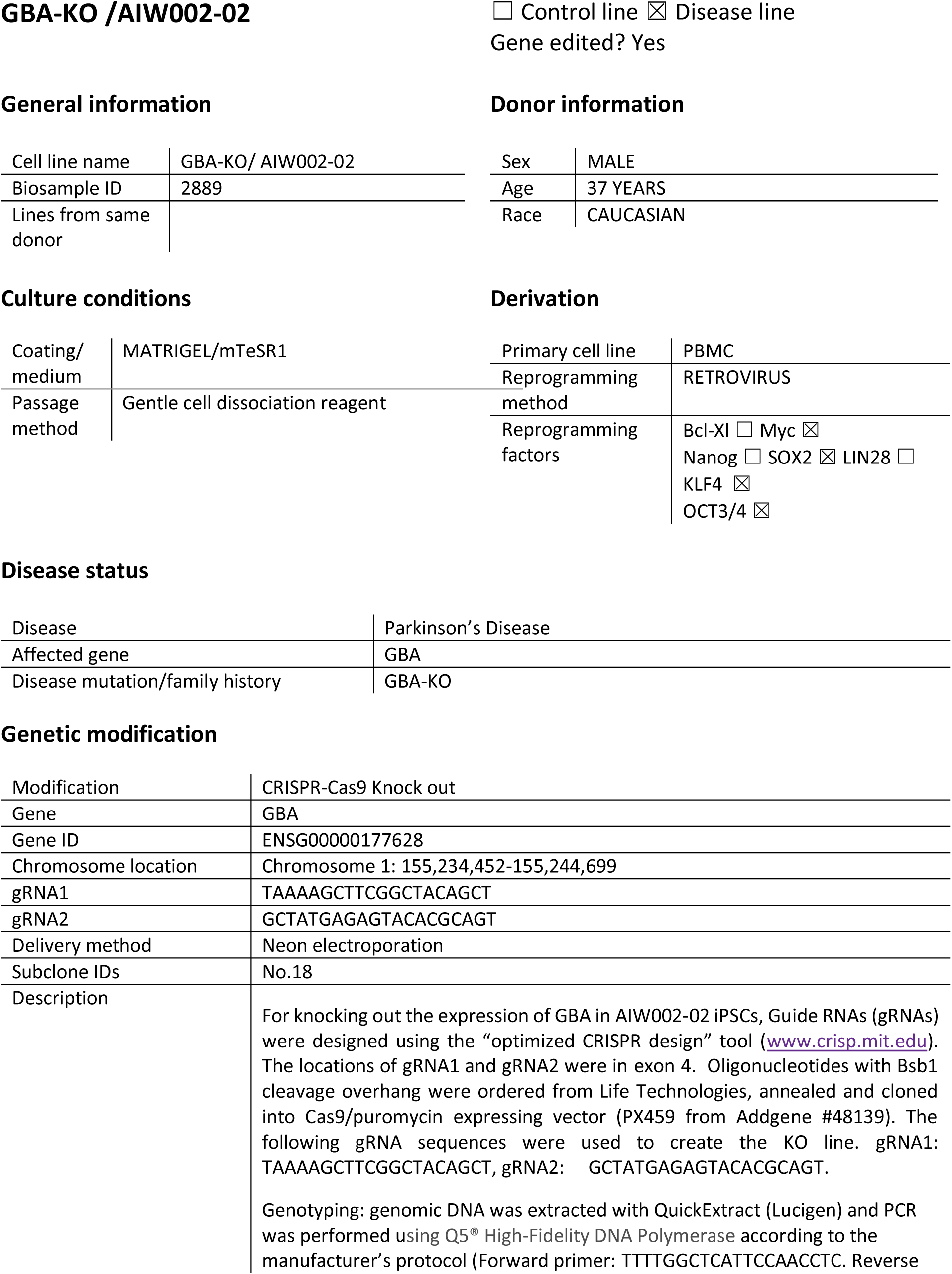
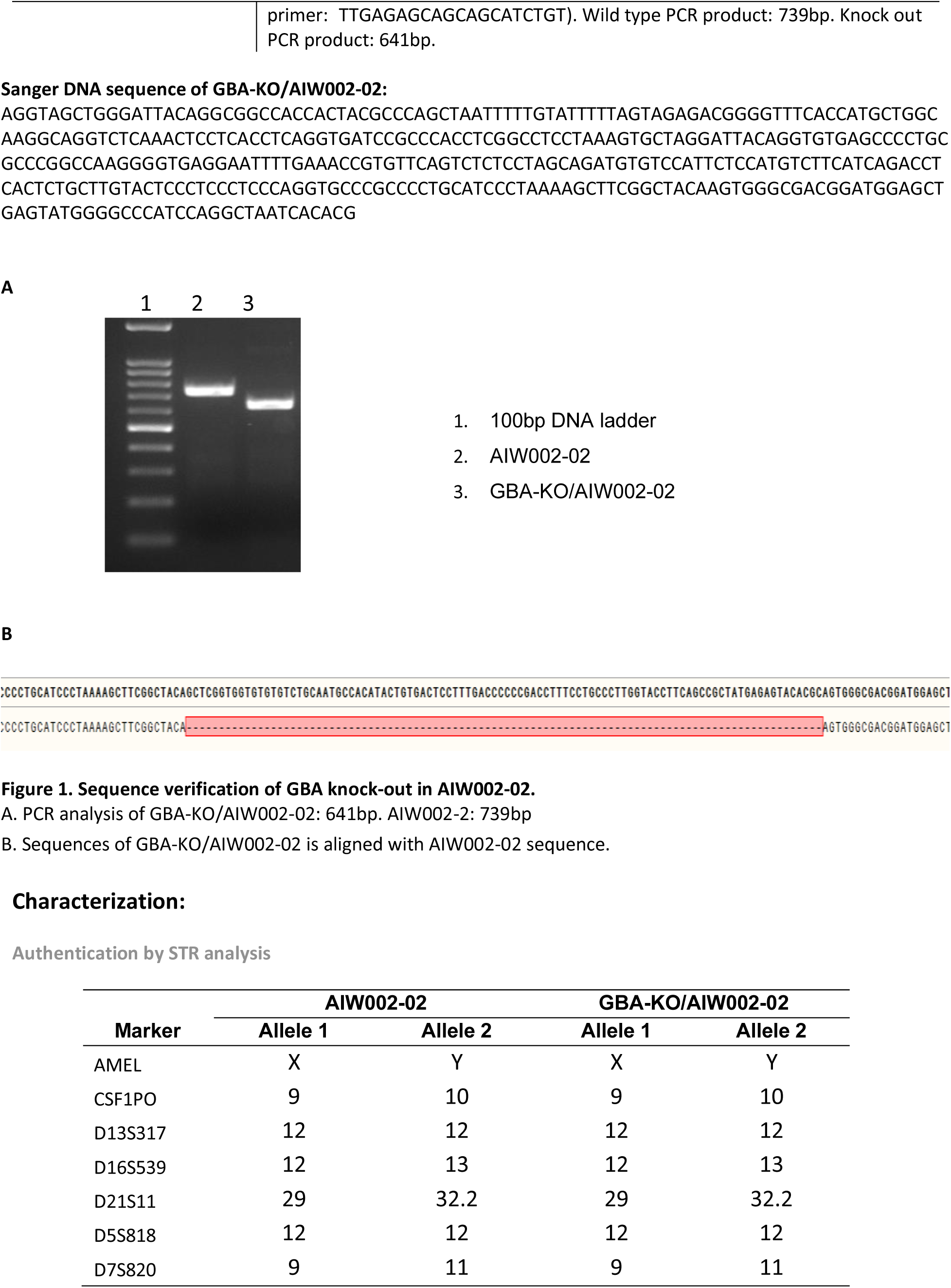
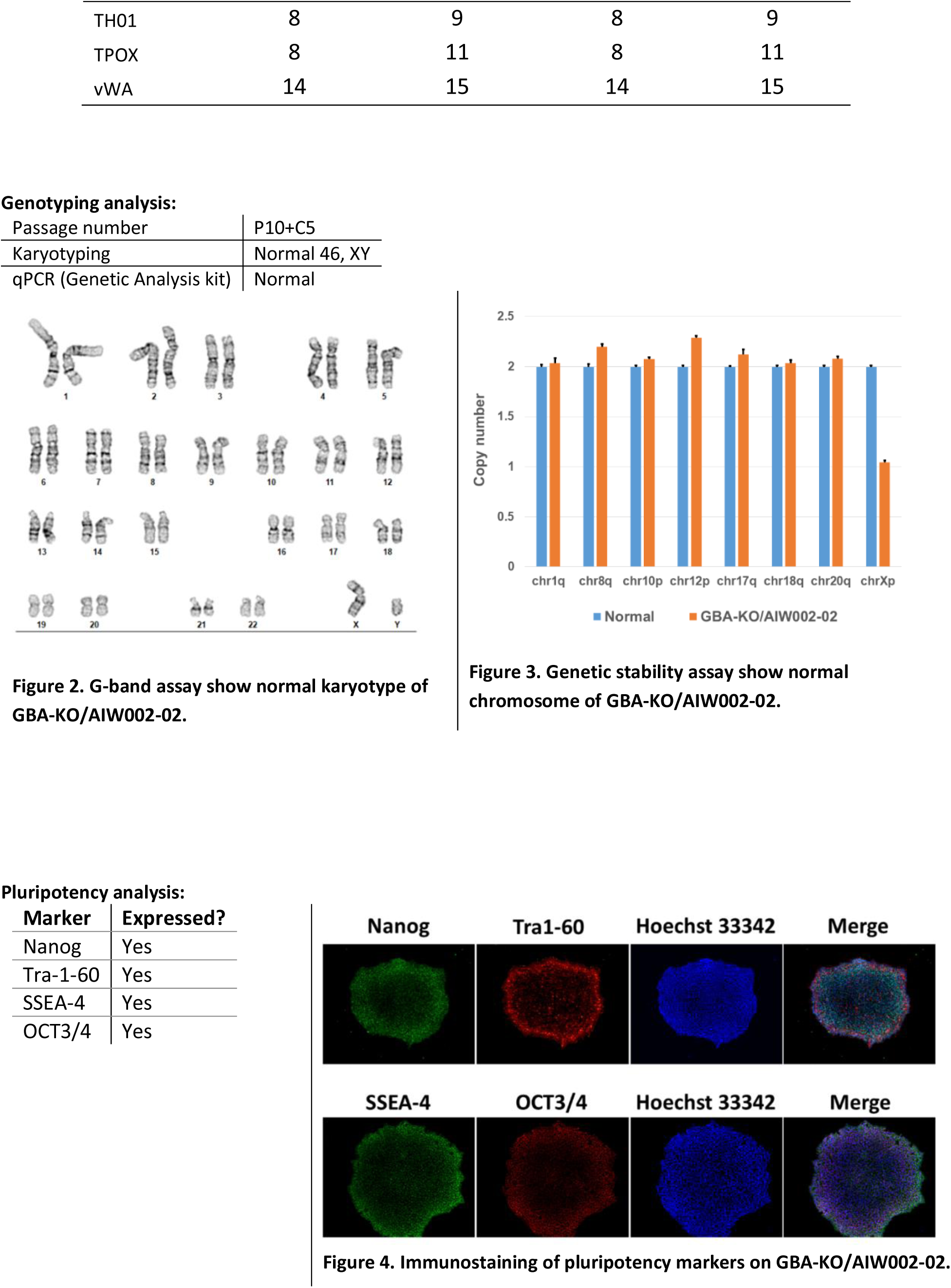
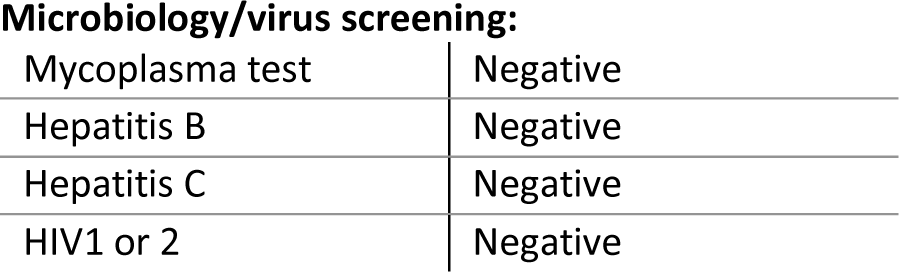

